# FLEXIBLE, PRODUCTION-SCALE, HUMAN WHOLE GENOME SEQUENCING ON A BENCHTOP SEQUENCER

**DOI:** 10.1101/2024.11.01.620975

**Authors:** Kevin Green, Benjamin Krajacich, Kelly Wiseman, Peter T. Mains, Samantha Robertson, Sophie Billings, Mitch Sudkamp, Ching Shing Lo, Bryan R. Lajoie, Semyon Kruglyak, Shawn Levy, Junhua Zhao

## Abstract

Human whole-genome sequencing (hWGS) provides comprehensive genomic information that can potentially help guide research in disease prevention and treatment and ultimately improve human health. Recent advancements in sequencing technology have improved sequencing quality and further reduced sequencing costs on bench-top sized instruments, making whole-genome sequencing an accessible technology for broader use. Here, we demonstrate the feasibility of a large whole genome sequencing project using a benchtop sequencer in a small laboratory setting, on a scale previously reserved for production-scale factory-sized machines. In this project, 807 samples were prepared and sequenced across 313 flow cells, with high sequencing quality at a median %Q30 of 96.6% and a median %Q40 of 89.31%. To screen library quality and maximize sample yield, we utilized 48-plex sample pre-pool ‘QC’ runs to provide >1x coverage per sample prior to sample pooling and full-depth sequencing. With this strategy, we consistently achieved >30x human whole genome sequencing of three-plex sample trios with standard settings or up to 4 samples per run with a high-throughput run setting. Additionally, this low-pass data provided valuable sample-level insights, allowing for detection of chromosome copy number variations (CNV) prior to full-depth sequencing. With three instruments running concurrently, >2,800 30x human whole genomes could be sequenced per year. To further demonstrate additional flexibility present in the platform, we also explored two different use cases 1) large insert sizes (1kb+) library to achieve superior genome coverage; 2) a proof of concept for rapid WGS sequencing to minimize sample to answer turnaround time for time-critical sequencing applications. Sequencing of a 2×100 >30x human WGS can be achieved in <12 hours and subsequent generation of fastq, bam and vcf in <1 additional hour. This study provides a cost-effective and flexible real-world demonstration of achieving both high quality hWGS sequencing and instrument flexibility without the need for complex batching schemes or factory-sized sequencers.

## INTRODUCTION

Whole-genome sequencing (WGS) is a pivotal tool in the research and study of rare diseases (Bagger et al., 2024; Bertoli-Avella et al., 2021; Stranneheim et al., 2021; Vinkšel et al., 2021). WGS can provide understanding for individuals with rare genetic disorders, offering molecular clarity crucial for research in targeted therapies and precision medicine (Brlek et al., 2024; Clark et al., 2018; Yadav et al., 2023). WGS is recognized as a suitable first-tier action for individuals with rare genetic disorders, potentially supplanting other research approaches such as chromosomal microarray analysis and whole-exome sequencing (Marshall et al., 2020; Souche et al., 2022). Through the analysis of rare alleles using WGS, it becomes feasible to uncover the presence of specific traits related to rare diseases in certain individuals (Turro et al., 2020). Additionally, rapid WGS has demonstrated the ability to provide insights into the underpinnings of genetic disorders within a remarkably short timeframe, potentially enabling prompt interventions and enhanced outcomes, particularly in critically ill infants (Farnaes et al., 2018; Lewis et al., 2023; Marom et al., 2024; Petrikin et al., 2018).

Previous studies have shown that Avidite Base Chemistry (ABC) on the Element Biosciences AVITI™ benchtop sequencer is directly compatible with existing upstream and downstream workflows and can be used to study rare genetic diseases associated with inherited retinal degeneration (IRD) phenotype and other rare, undiagnosed neurological disorders (Biswas et al., 2024; Ramsey et al., 2023). Additionally, the ABC chemistry on AVITI allows the use of WGS libraries with inserts longer than typical short-read sequencing libraries (>1kb compared to 300-400bp) to achieve higher mapping and variant calling accuracy compared to Illumina sequencing at the same coverage, with greater benefits at reduced coverage (Carroll et al., 2023).

This study presents use-cases and applications over a range of experimental designs that leverage the AVITI system. We describe the generation of 800+ WGS samples to >30x depth with optimizations used to improve reliability and reduce re-queueing of samples, representing high-throughput and scalable use. We describe the generation and sequencing of >1kb+ libraries shown to provide more accurate variant calling and detection, demonstrating beneficial effects on benchmarking and variant calling resolution and accuracy. Finally, we describe the use of AVITI to perform full coverage >30x depth of a human genome in 12.5 hours. Rapid time to results is a critical feature in many emerging applications including neonatal inherited disease, oncology, and infectious disease. These projects and applications demonstrate the flexibility in usage and ability to scale large throughput projects onto a benchtop system with consistent and reproducible results.

## MATERIALS AND METHODS

### Sample Handling and gDNA Quantification

To represent high-throughput and scalable use at two scales, Project 1 processed 807 unique samples while Project 2 processed 92 unique samples. All genomic DNA samples were transferred into 1 mL Matrix tubes within the Matrix tube rack (Cat: 3741, Thermo Fisher) in 96-well plate format and assigned unique gDNA sample barcode, provided by the bottom of each tube for sample tracking. To reduce QC cost, we miniaturized a dsDNA quantification assay based on the Quantifluor dsDNA system (Promega, USA). The 200 uL stock reaction volume was reduced to 40 uL total (36 uL working solution with 4 uL of diluted sample or standard) and samples were quantified in triplicate on the Agilent Neo2 plate reader using an excitation/emission ratio of 504/530. Samples received contained 10ng to 5ug of DNA per sample, with varying DNA Integrity number scores (DIN) ranging from 1.6-9.2 (Tapestation Genomic DNA Reagents, Agilent Technologies, USA).

### Library preparation

To generate libraries ranging from 350bp insert to >1kb, gDNA (500ng-1ug) was mechanically sheared in 55uL of 10 mM Tris using the Covaris ME220 instrument (Woburn, MA, USA) with sonication protocols adjusted to enrich gDNA fragment distribution for the desired insert size range (Table 1). Library conditions were optimized to produce aligned insert sizes of 350bp, 600bp, and >1kb. Table 1 outlines the shearing and SPRI ratios used during double sided size selection optimized for each library prep condition. To generate the insert sizes centered around 350 bp, optimal post-ligation double-sided SPRI clean up ratios were 0.46x/0.62x. To produce libraries with an aligned insert size centering around 600 bp, mechanical shearing time was increased to 30 seconds per sample with a reduction in the number of cycles per burst, resulting in larger sheared gDNA fragments. Additionally, the double-sided SPRI cleanup ratios were adjusted to 0.4x/0.52x to select for larger libraries after post-ligation cleanup. To generate the longest insert libraries, >1kb in length, mechanical shearing time was reduced from 30 to 5 seconds prior to library prep to ensure longer gDNA fragments were present. After adapter ligation, two 0.8x post adapter ligation cleanups were completed to remove all small library fragments and adapter dimers, followed by a double-sided SPRI cleanup with ratios as 0.3x/0.42x (Table 1). It is imperative to remove all small library material during the long insert preparation to ensure efficient sequencing of only the longest library fragments. Presence of smaller library molecules will preferentially amplify during sequencing, resulting in smaller aligned insert sizes than desired. Samples which did not meet input requirements for PCR Free library prep (> 500 ng) underwent PCR amplification after adapter ligation and size selection. To ensure sufficient material was available for PCR amplification, a single sided size selection using 0.6x SPRI ratio was used after the 0.8x post-ligation SPRI cleanup. This single sided selected library was amplified according to manufacturer’s instructions and provided master mix and primers. Libraries were quantified via Promega QuantiFluor assay or Qubit (Thermo), and quality checked using the Agilent TapeStation High Sensitivity D5000 reagents.

**Table 1:** Mechanical shearing and SPRI conditions used in library preparation.

| Covaris Shearing |  |  |  |  |  |  | SPRI Ratio |
| --- | --- | --- | --- | --- | --- | --- | --- |
| Insert Size | Duration (s) | Peak Power | Duty % Factor | Cycles/Burst | Avg. Power | Iteration | Double Sided Size Selection |
| 350 bp | 10 | 50 | 20 | 1000 | 10 | 7 | 0.46x/0.62x |
| 600 bp | 30 | 50 | 20 | 200 | 10 | N/A | 0.4x/0.52x |
| > 1kb | 5 | 50 | 20 | 200 | 10 | N/A | 0.3x/0.42x |

Higher throughput sample processing (intake QC, library preparation and QC, and circularization and QC) and sequencing by a single operator utilizing benchtop instruments is easily accomplished as outlined in Figure 1. Project 1 processed 807 unique patient samples using the KAPA HyperPrep Library Prep kit (Roche, USA). The first 77 samples were processed using the 350 bp insert size conditions outlined in Table 1, then shifting to the 600 bp conditions for the remaining 730 samples. Project 2 processed 92 unique samples using the >1kb conditions using the Element Elevate Library Prep workflow (Element Biosciences, USA) with Element Full-lengths UDIs.

**Figure 1:**
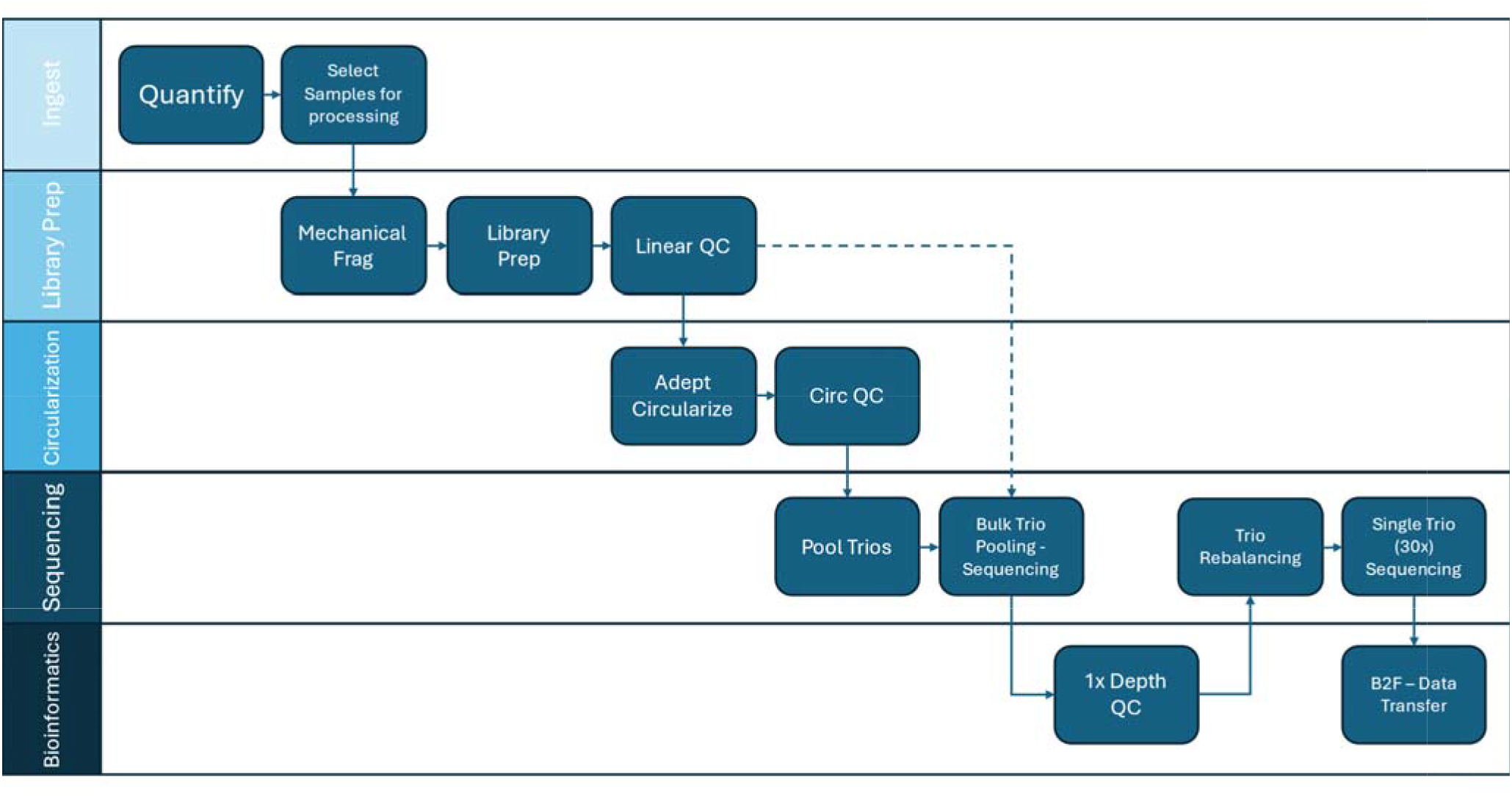
Laboratory workflow from sample ingest to data analysis.

All projects utilized the Rapid Adept ™ Workflow with NaOH Denaturation protocol (Element Biosciences, USA) to generate circular libraries unless otherwise stated. The final circularized libraries were quantified using the Applied Biosystems SYBR Green Universal PCR Master mix (Thermo Fisher Scientific, USA) and primers provided within the Rapid Adept kit. Circular libraries were diluted to 1 nM, then used as input into NaOH denaturation at the final library loading concentration along with PhiX libraries as a sequencing control.

### Sequencing

To understand the impact of insert size on benchmarking, HG001 gDNA (Coriell) was used as input material and libraries were prepared with the 350 bp, 600 bp, and >1kb insert workflows as outlined above in Table 1. Libraries were sequenced to 35x coverage with 2×150 cartridges. Large insert libraries were sequenced using an optimized recipe with longer amplification time to improve the intensity (recipe available upon request).

To effectively sequence Project 1, a “Bulk QC” sequencing run of up to 16 trios (48 individual samples) were pooled equimolar before being loaded onto a single AVITI Cloudbreak™ flow cell and sequenced for 300 cycles (2×150) with indexing reads. Using this strategy, 48 human WGS samples can be sequenced to > 1-2x coverage for initial secondary QC to scan for library preparation biases prior to full depth sequencing runs. Based on “Bulk QC” inter-trio balance of individual indexes, each trio was rebalanced based on read yield to increase the evenness of the indices before sequencing at full depth across one flow cell per trio. The rebalanced trios were then sequenced using 2×150bp cartridges.

The sequencing strategy for Project 2 was similar to Project 1. After library prep, samples were pooled for Bulk QC runs to complete 1x QC then pooled into sets of 4 samples for individual 2×150 sequencing runs using Cloudbreak chemistry on the AVITI instrument. Large insert libraries were sequenced with paired-end, 2×150 sequencing using an optimized recipe with longer amplification time to improve the intensity (recipe available upon request). Full depth flow cells were loaded to a target density of 600-700M polonies per side of the AVITI instrument to maximize quality at the larger insert size.

To demonstrate minimal time from sample to results, two rapid WGS (rWGS) sequencing strategies were developed. Strategy 1 – Two flow cells within 12 hours, was performed by sequencing a single circular library across two flow cells on a single AVITI instrument with a custom recipe to reduce sequencing time. Both runs were set up simultaneously for 2×100 low output AVITI Kit targeting 250M reads per flow cell and completed within 12 hours. Strategy 2 – one flow cell for 24 hours, utilized a single flow cell and the Cloudbreak Freestyle™ compatibility chemistry, allowing for direct loading of a linear library into the AVITI sequencer and sequencing 2×150 cycles in 24 hrs with mid-output scanning targeting 500 M reads per flow cell.

### Secondary analysis

FASTQ files were generated via Bases2Fastq and used as input into analysis for down sampling and additional alignment. (https://docs.elembio.io/docs/bases2fastq/introduction/). For Project 1 and 2 Bulk QC, FASTQ files were down sampled to 1x coverage for alignment to the hg38 human reference and analysis with the Sentieon DNAScope pipeline (https://www.sentieon.com/). Insert size, GC/AT bias, and uniformity were calculated per sample and tracked across sample prep batches. For benchmarking and analysis, input FASTQs were down sampled to 35X and aligned using Sentieon bwa unless otherwise noted. Benchmarking was accomplished using Google DeepVariant 1.6 and NISTv4.2.1 truth vcf/bed. Sample handling for large scale projects was monitored using metadata/chrX coverage and NGS fingerprinting tools in Picard Tools (Broad Institute, https://gatk.broadinstitute.org/hc/en-us/articles/360041696232-Detecting-sample-swaps-with-Picard-tools) to monitor potential sample swaps that may have occurred during sample processing.

## RESULTS

### Enhanced Benchmarking Performance with Longer Insert Size Libraries

Based on simulation data (https://www.elementbiosciences.com/resources/the-impact-of-insert-length-on-variant-calling-quality-in-whole-genome-sequencing), libraries containing a larger insert size could improve mapping and coverage of difficult regions in the genome, even if sequenced within the short reads setting (300 cycles). In this project, we optimized library prep and sequencing recipes to enable sequencing of larger insert size libraries to generate high quality sequencing data for benchmarking. Table 1 outlines the optimized library prep conditions to generate PCR Free, high-quality WGS libraries at the 350 bp, 600 bp, and >1kb insert sizes. Median aligned insert sizes of the final, sequenced libraries were 386 bp (SD 122.46 bp), 584 bp (SD 170.10 bp) and 1339 bp (SD 444.26 bp) for the small, medium, and large inserts respectively (Figure 2). Run quality and alignment performance are summarized in Table 2.

**Table 2:** Sequencing run quality across optimized library sizes when sequenced 2×150 using Cloudbreak chemistry.

| Library Size | Mean Base Quality | Percent Q30 | Percent Aligned to Reference |
| --- | --- | --- | --- |
| 350 bp | 42.85 | 96.57 | 99.88 |
| 600 bp | 40.74 | 92.88 | 99.82 |
| >1kb | 41.33 | 93.35 | 99.80 |

**Figure 2:**
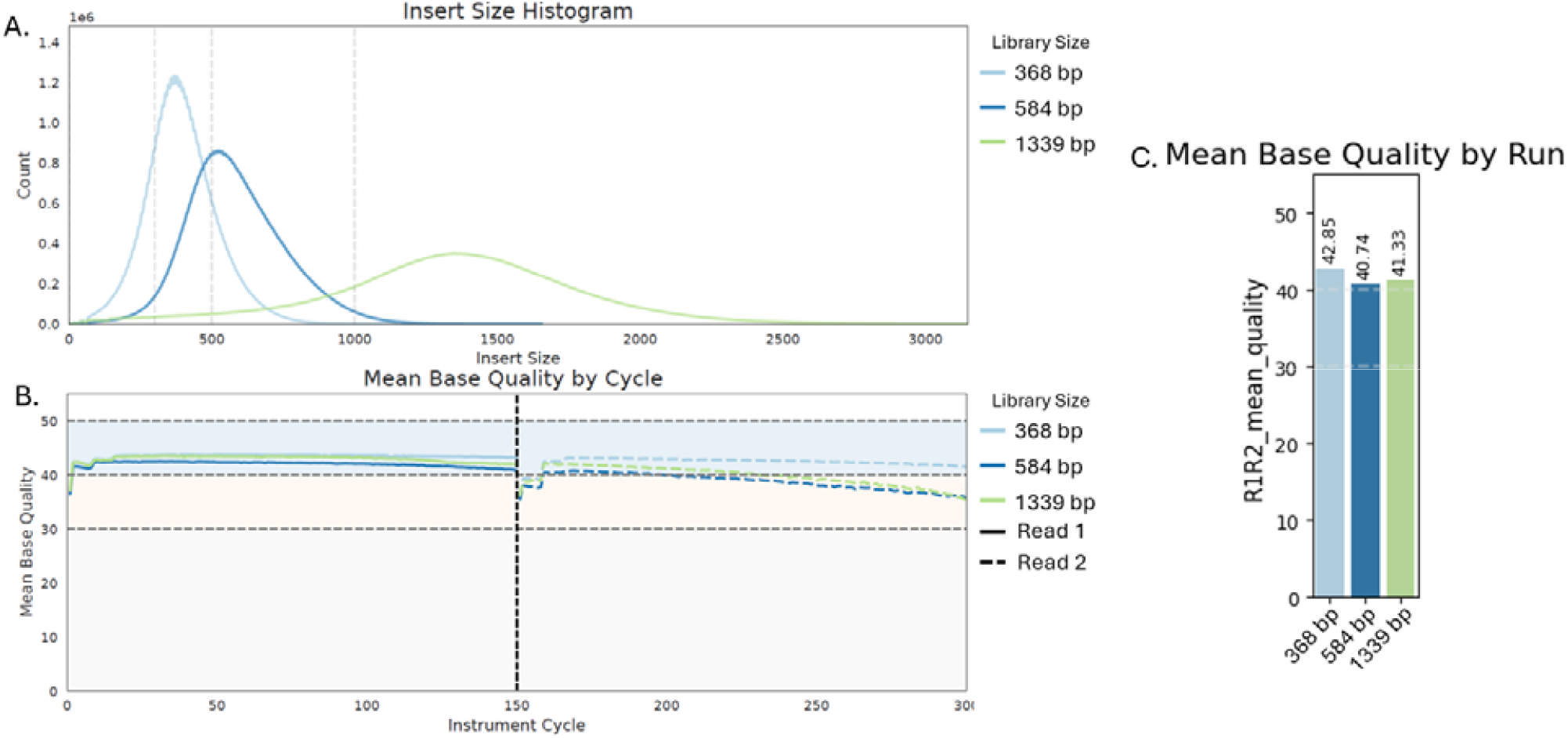
A) Median aligned insert size of the small (386 bp, SD 122.64 bp), medium (584 bp, SD 170.10 bp), and large (1339 bp, SD 444.26 bp) insert libraries. Insert size is calculated by alignment to HG001 reference genome. B) Base quality remained high across all cycles of the run. C) Mean base quality by run for each insert size.

To compare benchmarking results across multiple runs, each library was down sampled to ∼360 M reads for analysis, resulting in 35x coverage across the genome. Benchmarking of HG001 improved as insert size increased with total benchmarking errors reduced from 23044 in samples with the 350 bp insert size to 17112 with 1300 bp insert size, a reduction of 25.7%. Nearly all the improvement originated from a reduction in SNP False Negatives (FN) which reduced by 36.9% from 14827 errors in the smallest library (386 bp) to 9348 errors in the largest library (1339 bp) (Figure 3). The larger insert sizes contain fewer errors indicating that large insert size fragments provide better coverage across difficult regions within the human genome (Figure 4). Benchmarking F1 scores for both SNPs and INDELs across all regions increased as insert size increased, from 0.9963 INDEL and 0.9970 SNP at the smallest insert size to 0.9966 INDEL and 0.9979 SNP for the > 1kb insert size library. SNP F1 scores in the difficult regions (such as contexts with homopolymers and tandem repeats, etc.) drove this overall F1 score increase with the smaller insert size having an SNP F1 score of 0.9846, and the > 1kb insert size yielded an SNP F1 score of 0.9897.

**Figure 3:**
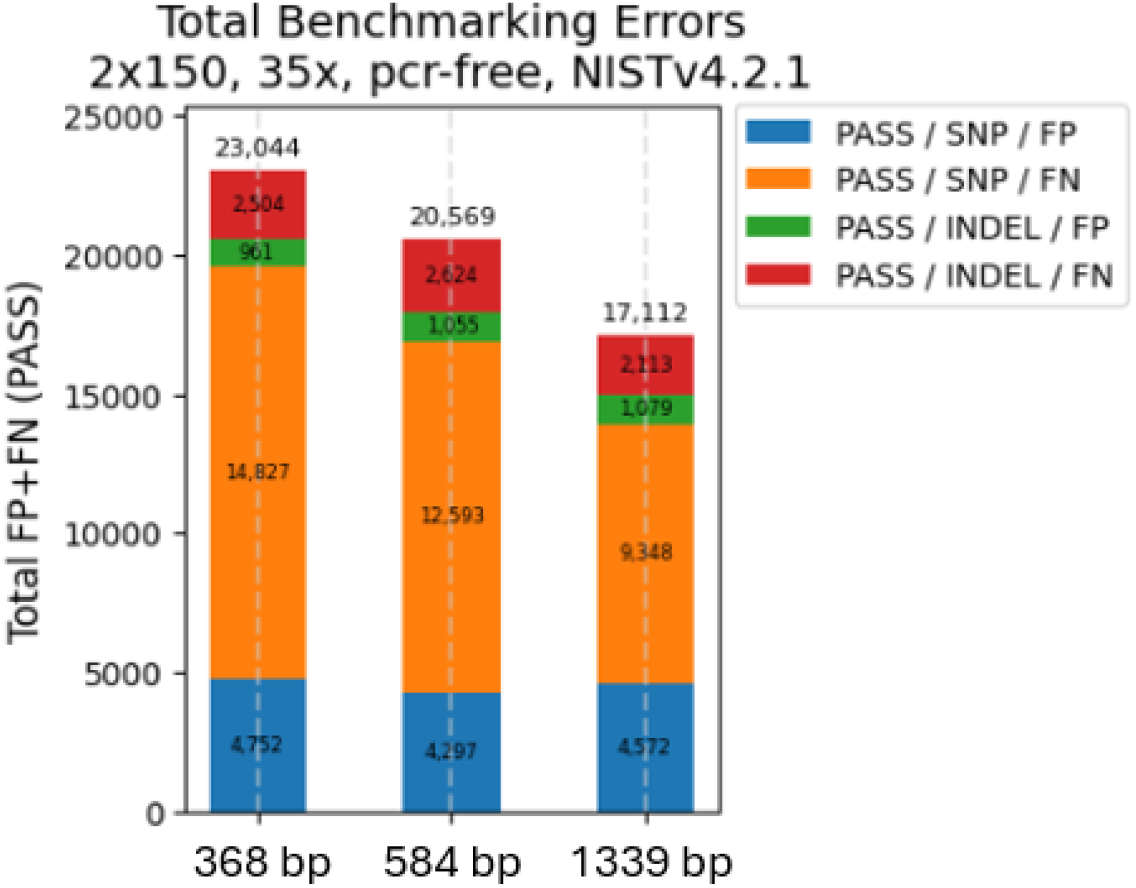
Increase in library insert size reduces total error during variant calling.

**Figure 4:**
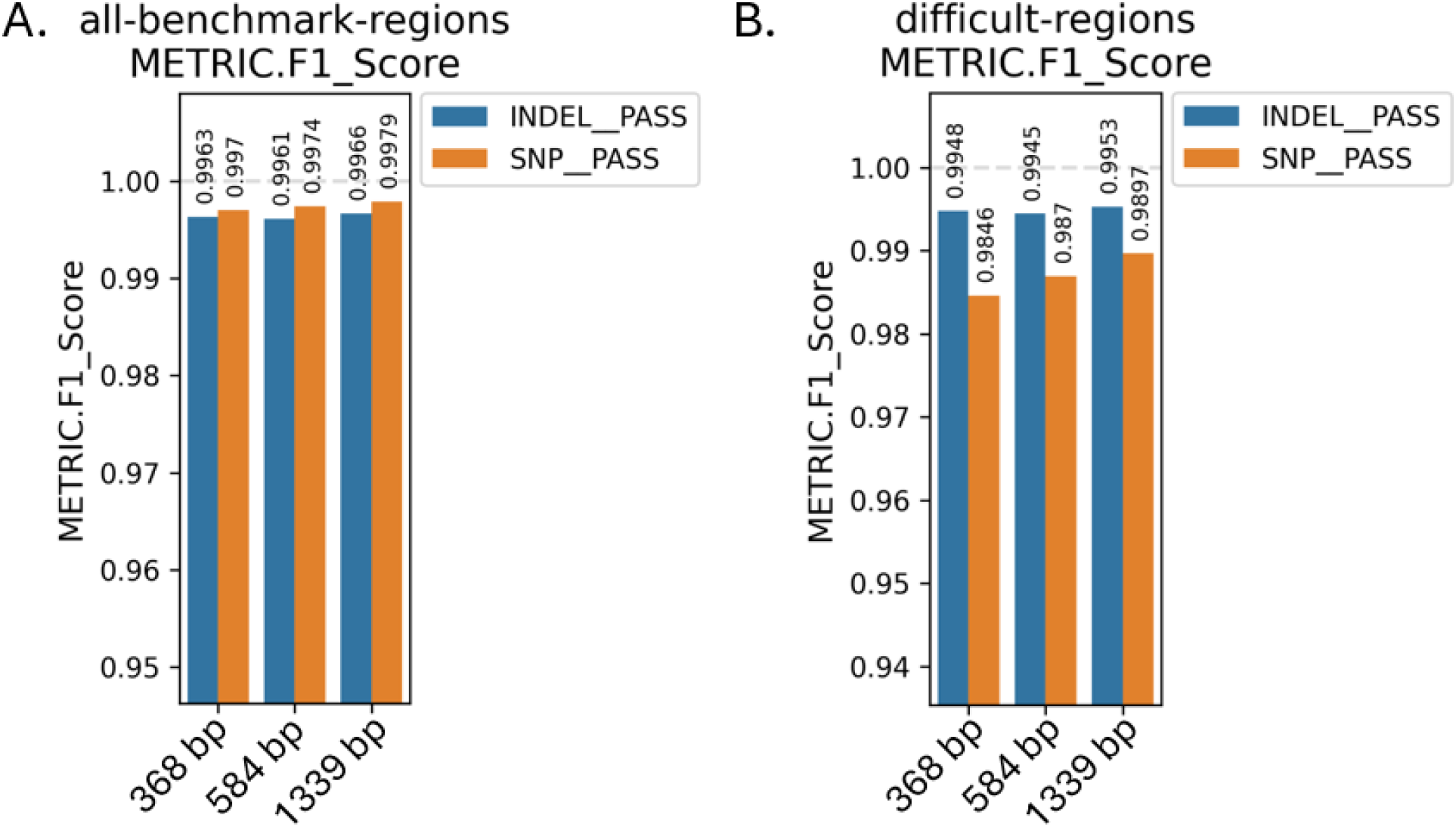
A) Benchmarking F1 scores for SNPs and INDELs across all regions of the genome of the 3 insert size libraries B) Benchmarking F1 scores for SNPs and INDELs of difficult regions of the genome.

### Achieving Reliable and Optimized Outcomes in Library Preparation

The two larger insert library prep methods (600 bp and >1 kb) were applied to both Projects 1 and 2 to leverage the benefit observed from larger insert sizes during benchmarking. To complete Project 1, 880 libraries were prepared PCR Free across 17 batches to fully process the 807 samples provided. Due to low quality or confirmatory testing needing to be completed, 73 samples needed to be reprepared starting from gDNA. Of the 880 libraries generated to complete the project, 775 libraries were processed as PCR Free preparations, while 105 samples required additional amplification during the library preparation workflow due to low input gDNA. 698/775 PCR Free libraries were generated using the 600bp insert conditions, with the first 77 samples of the project being generated using the 350bp conditions as outlined in Table 1. While library size varied slightly between batch, the median aligned insert size across all samples processed with the 600bp insert conditions was 587 bp (n=698, SD=93.32, %CV=15.89%). The 105 samples prepared with a final PCR amplification (PCR Plus protocol) had a median aligned insert size of 432bp (n=698, SD=53.69, %CV=12.66%) (Figure 5).

**Figure 5:**
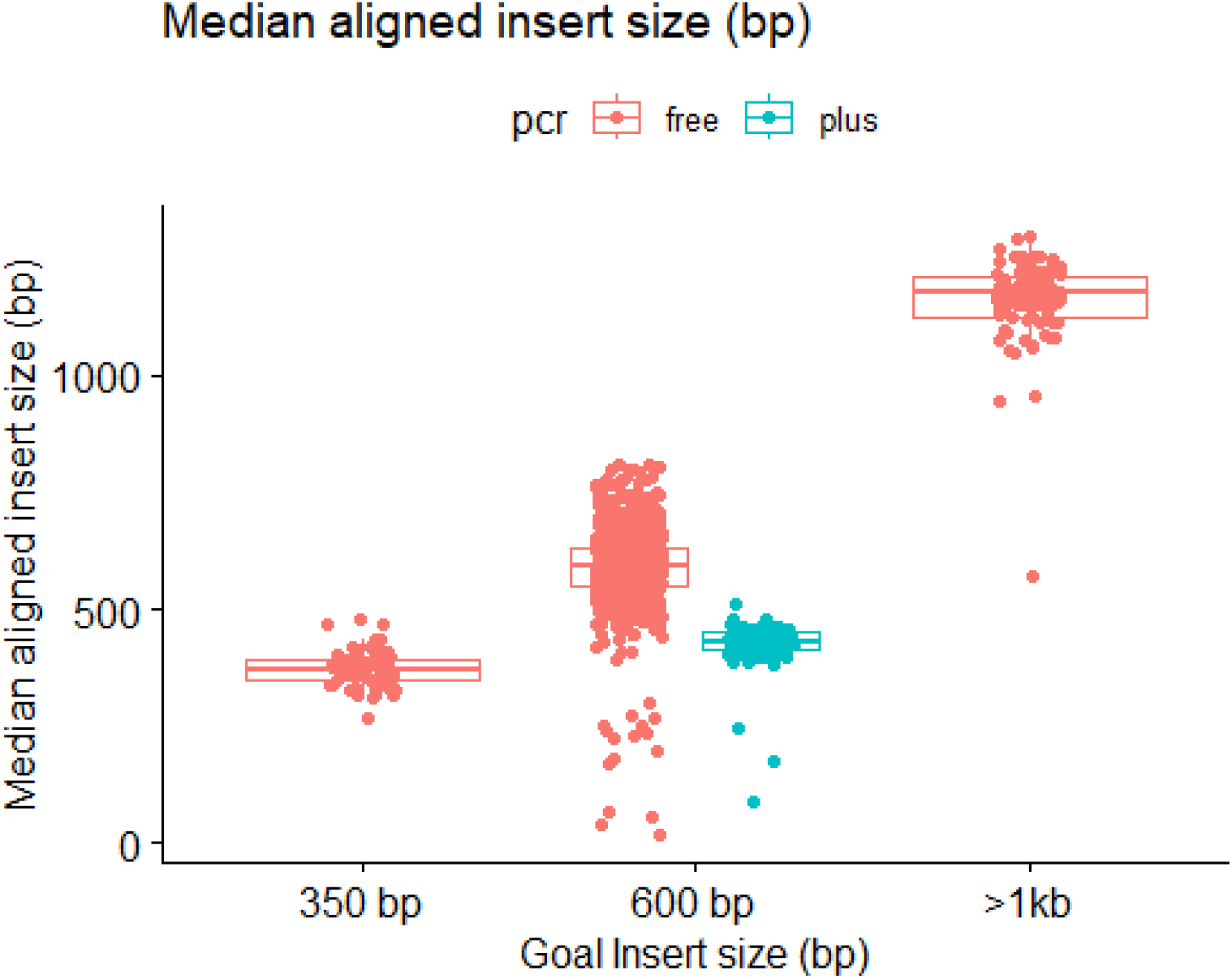
Median aligned insert size of PCR Free and PCR Plus libraries generated for Projects 1 and 2. The first 77 libraries for Project 1 were generated using the 350bp insert method, with the remaining samples being prepare using the 600bp insert or PCR Plus method. The libraries with small aligned insert sizes indicate potential microbial contamination.

Libraries for Project 2 were generated using the long insert conditions and generated libraries with an insert of 1-1.5 kb, 92 samples were prepared and sequenced using the Bulk QC method over 2 runs. Median aligned insert size across 92 samples averaged 1162 bp (n=92, SD=89.2, %CV=7.68) (Figure 2).

### Enhancing Library Loading Consistency for Optimal Sequencing Utilization and Coverage Depth

Accurate normalization of samples ensures the number of sequencing runs needed to complete each project are minimized, with improper normalization of samples potentially resulting in multiple costly and time-consuming resequencing events for larger scale projects. To minimize the pooling variation in a cost-effective way, sample trios were pooled and sequenced at higher plexity for a “Bulk QC” to evaluate index balancing. The trios were then rebalanced after the Bulk QC run to bring the representation of each sample within the trio to ∼33%. Table 3illustrates an example Bulk QC and rebalance of 15 trio pools from Project 1. 45 individual samples were pooled into 15 trio pools and subsequently pooled equimolar into a single bulk pool for Bulk QC sequencing. Across the 15 pools, the average intra-trio index CV (CV of the 3 samples within a single pool) was 12.27% with a maximum intra-trio index CV of 47.55% of a single pool (n=15, SD=10.44%). After rebalancing the pools to ensure more equal representation of indexes within a trio based on the Bulk QC index representation data, the same 15 trios were sequenced to full depth on individual full sequencing runs and index representation calculated using index assignment. The rebalancing of the trios improved the index CV, reducing the average intra trio index CV to 4.41% with a maximum intra-trio index CV of 9.33% (n=15, SD=1.82%) (Table 3).

**Table 3:** Bulk QC inter trio CV calculated from the Bulk QC run (left column) and the final inter trio CV from the full depth sequencing run of each rebalanced trio (right column).

| Pool | Bulk QC trio CV (%) | Rebalanced CV (%) |
| --- | --- | --- |
| 1 | 47.55 | 3.26 |
| 2 | 8.36 | 4.34 |
| 3 | 4.79 | 3.02 |
| 4 | 11.53 | 3.31 |
| 5 | 7.50 | 6.69 |
| 6 | 8.79 | 3.36 |
| 7 | 19.09 | 5.59 |
| 8 | 12.79 | 5.09 |
| 9 | 10.36 | 2.90 |
| 10 | 1.82 | 2.17 |
| 11 | 15.16 | 3.87 |
| 12 | 10.25 | 9.33 |
| 13 | 2.00 | 3.03 |
| 14 | 14.00 | 6.24 |
| 15 | 10.10 | 3.92 |
| Average | 12.27 | 4.41 |
| SD | 10.45 | 1.82 |

Along with index representation, the Bulk QC runs provide valuable information on library preparation success. Success was quantified by measuring the genomic coverage variation at 1x coverage, providing information on how evenly each sample was sequenced across the genome. A genomic coverage variation metric of 0.1 (10%) was used as the maximum allowable variation for a given sample. Figure 6A illustrates sample genomic coverage variability across a subset of samples. Samples below the 0.1 coverage variation threshold displayed a more even and “flat” coverage (Figure 6B), while samples greater than 0.1 showed increased genomic variability and non-uniform coverage plot (Figure 6C).

**Figure 6:**
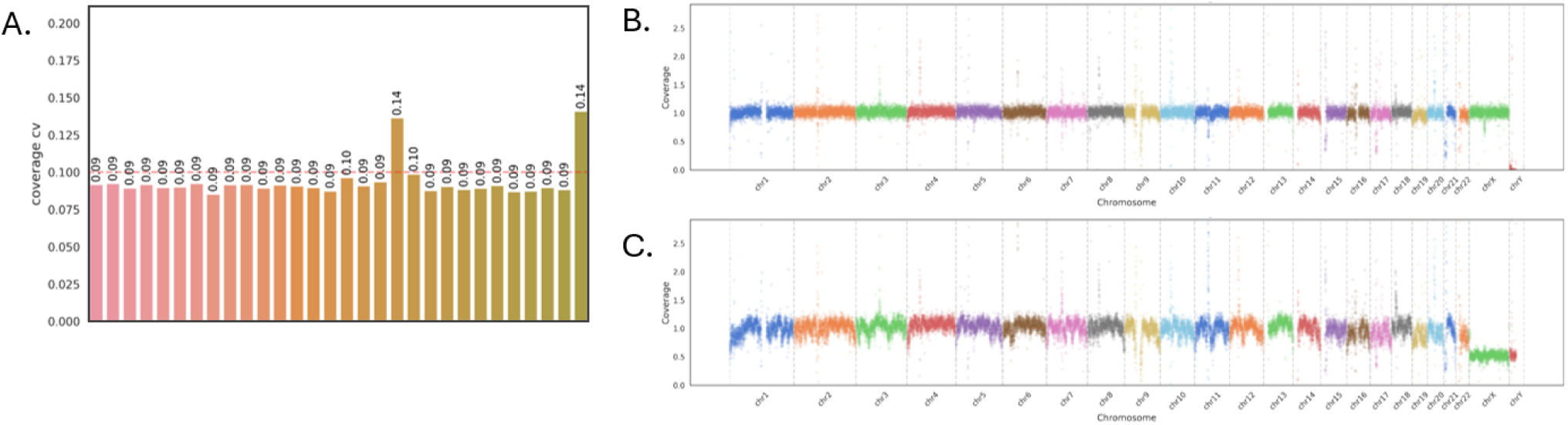
A) Genomic coverage CV at 1x coverage of 33 example samples. Samples with a CV greater than 0. were marked for additional screening and QC. B) Example genomic coverage CV plot at 1x of a sample with a CV of 0.09 yielding a “flat” coverage profile. C) Non uniform genomic coverage profile of a sample with a CV of 0.14.

### Assessing Sequencing Fidelity and Consistency

Sequencing quality across the large-scale >300 sequencing run Project 1 and the smaller >60 sequencing run long insert Project 2 remained high across all runs. 315 runs were needed to fully sequence the 807 samples to at least 35x coverage in Project 1. Across the 315 runs, median polony output was 978.1 M (n=315, SD=172.9 M) (Table 4, Figure 7) and median Q30 base quality was 95.5% (n=315, SD=5.2) (Table 4, Figure 8). Run success rate was very high with 313/315 (99.36%) runs successfully sequenced without failure. The two failed runs root cause were operator error. Project 2 applied a long-insert sequencing recipe and targeted a lower density output. It required 63 sequencing runs to fully sequence all 92 long insert libraries to full depth with a median Q30 of 94.73% (n=63, SD=1.86) (Table 4, Figure 7) and median polony output of 630.1 M (n=63, SD=128.1 M) (Table 4, Figure 8). To regularly check sequencing quality and consistency, baseline control libraries were sequenced as standard practice across multiple instruments during the completion of both Project 1 and Project 2. Five example baseline sequencing runs of the same control library over 6 months (Table 5)_were analyzed to compare Ti:Tv ratios and the precision and accuracy of SNP/INDEL variant calling using DeepVariant (Table 6). The SNP True Positives (TP) for all five baseline runs were compared pairwise and showed high concordance of called SNP TPs, with an average of 99.87% shared SNP TPs being called across all 5 runs. (Table 7).

**Table 4:**
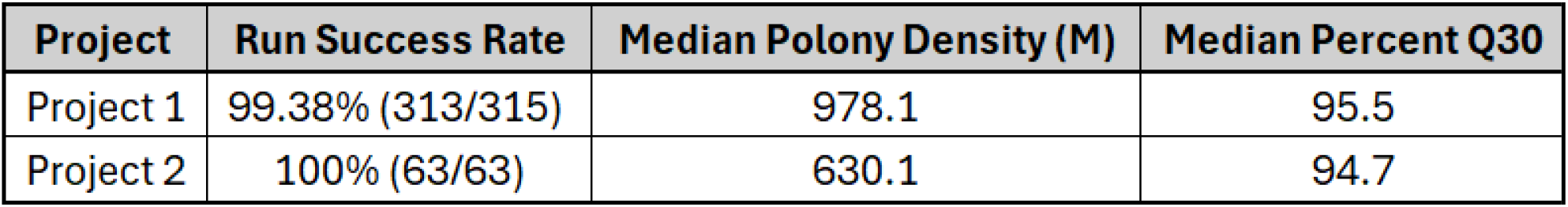
Project 1 and 2 sequencing run success rate and primary metrics.

| Project | Run Success Rate | Median Polony Density (M) | Median Percent Q30 |
| --- | --- | --- | --- |
| Project 1 | 99.38% (313/315) | 978.1 | 95.5 |
| Project 2 | 100% (63/63) | 630.1 | 94.7 |

**Table 5:** Primary metrics of the 5 example baseline sequencing runs analyzed for concordance of SNP true positives.

| Metric | Baseline 1 | Baseline 2 | Baseline 3 | Baseline 4 | Baseline 5 |
| --- | --- | --- | --- | --- | --- |
| Run Date | 7-Sep-23 | 25-Oct-23 | 12-Dec-23 | 20-Jan-24 | 3-Feb-24 |
| Run configuration | 2x150 | 2x150 | 2x150 | 2x150 | 2x150 |
| Reads Passing Filter (M) | 1042.3 | 1091.5 | 909.1 | 1051 | 950.7 |
| Mean Base Quality | 41.92 | 41.06 | 41.96 | 42.24 | 42.56 |
| Bases >Q30 | 94.62 | 92.99 | 94.21 | 95.2 | 95.5 |

**Table 6:**
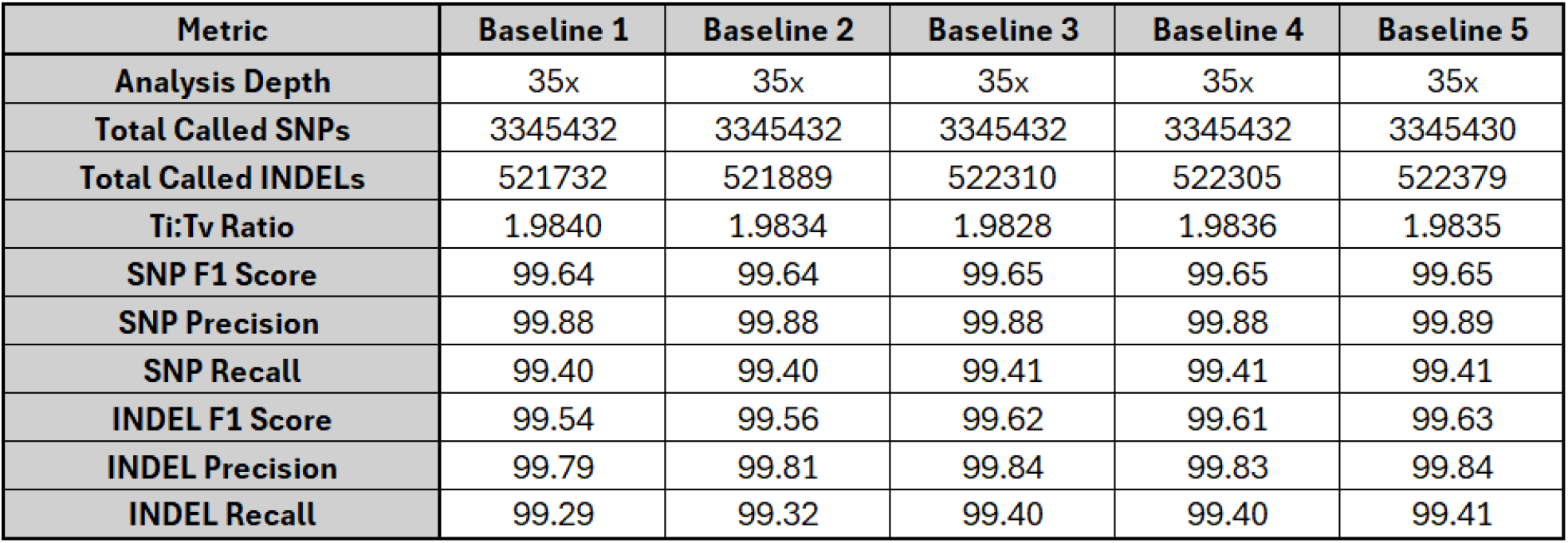
Baseline libraries secondary analysis comparison. HG002 down sampled to 35x depth and the Ti:Tv ratios, F1 score, recall, and precision of SNPs and INDELs was calculated.

| Metric | Baseline 1 | Baseline 2 | Baseline 3 | Baseline 4 | Baseline 5 |
| --- | --- | --- | --- | --- | --- |
| Analysis Depth | 35x | 35x | 35x | 35x | 35x |
| Total Called SNPs | 3345432 | 3345432 | 3345432 | 3345432 | 3345430 |
| Total Called INDELs | 521732 | 521889 | 522310 | 522305 | 522379 |
| Ti:Tv Ratio | 1.9840 | 1.9834 | 1.9828 | 1.9836 | 1.9835 |
| SNP F1 Score | 99.64 | 99.64 | 99.65 | 99.65 | 99.65 |
| SNP Precision | 99.88 | 99.88 | 99.88 | 99.88 | 99.89 |
| SNP Recall | 99.40 | 99.40 | 99.41 | 99.41 | 99.41 |
| INDEL F1 Score | 99.54 | 99.56 | 99.62 | 99.61 | 99.63 |
| INDEL Precision | 99.79 | 99.81 | 99.84 | 99.83 | 99.84 |
| INDEL Recall | 99.29 | 99.32 | 99.40 | 99.40 | 99.41 |

**Table 7:** Pairwise comparison of 5 example baseline runs analyzing percent shared SNP true positives between eac run using the same HG002 control library.

| % Shared SNP True Positives |  |  |  |  |  |
| --- | --- | --- | --- | --- | --- |
| Baseline 1 | Baseline 2 | Baseline 3 | Baseline 4 | Baseline 5 | Run |
|  | 99.86% | 99.87% | 99.87% | 99.87% | Baseline 1 |
|  |  | 99.87% | 99.87% | 99.87% | Baseline 2 |
|  |  |  | 99.88% | 99.88% | Baseline 3 |
|  |  |  |  | 99.88% | Baseline 4 |
|  |  |  |  |  | Baseline 5 |

**Figure 7:**
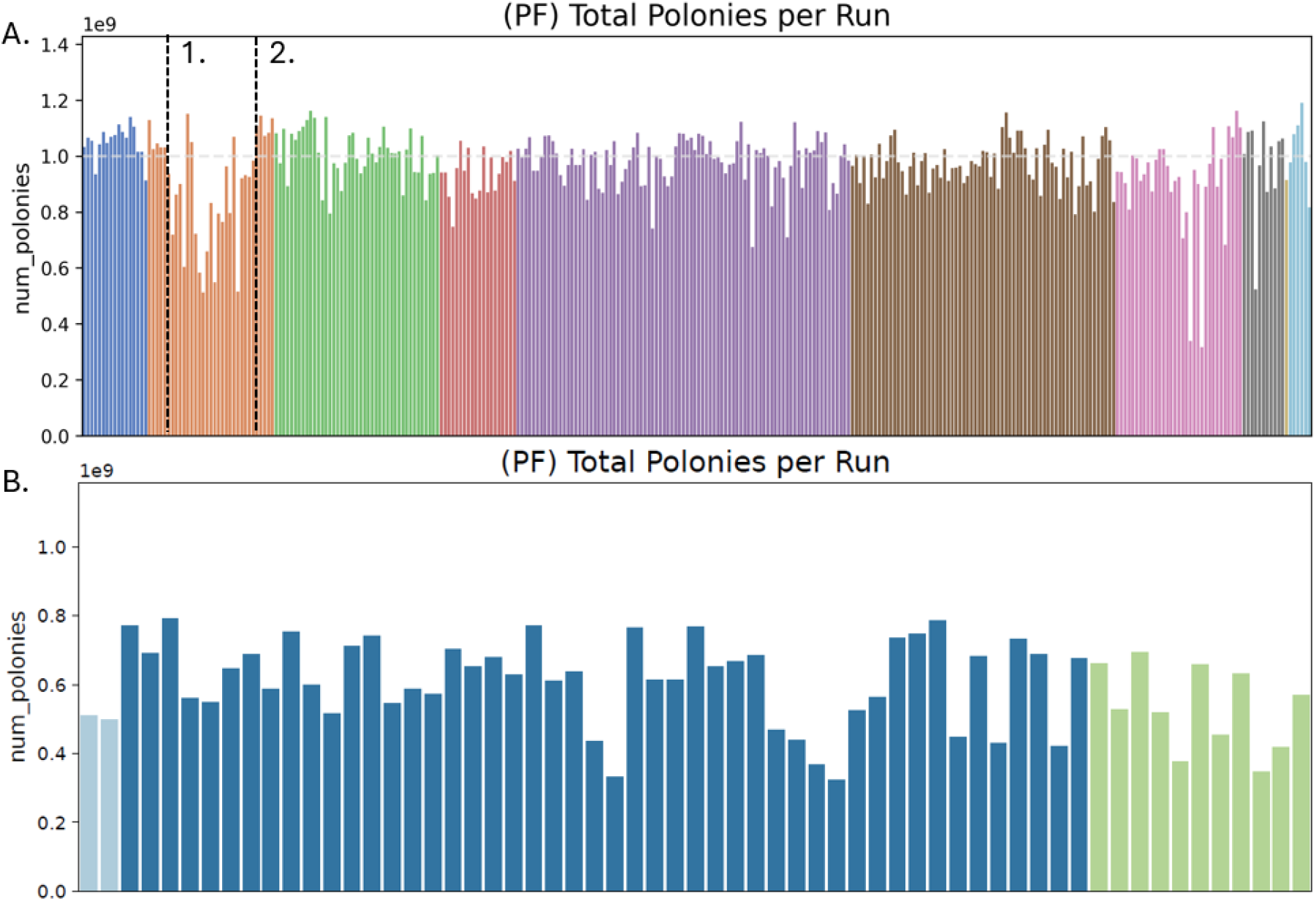
Total polonies per run over time of A) 600bp insert size Project 1 and B) >1kb+ insert library Project 2.

**Figure 8:**
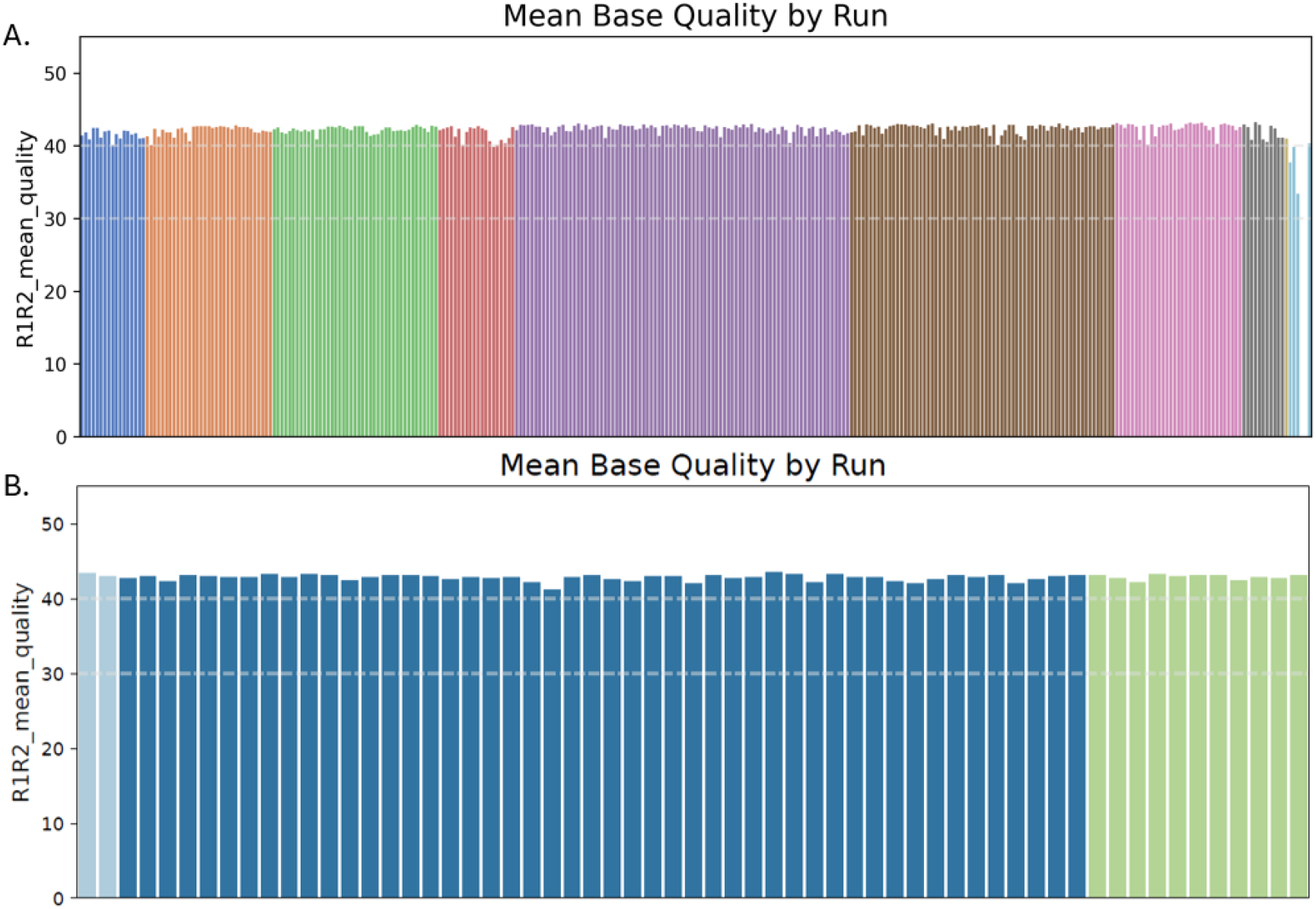
Mean base quality by run over time of A) 600bp insert size Project 1 and B) >1kb+ insert library Project 2. Dashed horizontal lines indicate Q40 and Q30 quality scores.

### rWGS Strategies: High Quality Sequencing and Accurate Variant Calling within 12 hours

In certain applications and use cases, sequencing and data analysis turnaround time (TAT) is prioritized over sample number throughput and batching. This is particularly true when sequencing is used in research studies for neonates and infants who potentially suffer from rare genetic diseases (Farnaes et al., 2018; Kingsmore et al., 2024; Lunke et al., 2023). Rapid Whole Genome Sequencing (rWGS) strategies can be employed to generate high quality data in as little 12 hours. To reduce overall sequencing time to fit within 12 and 24 hours, per cycle optimization of flow rates, contact times, and washes were reduced. Two rWGS strategies were employed to generate high quality data. Strategy 1 utilized two flow cells on both sides of a single instrument, allowing each side to quickly generate data and be combined for analysis in 12 hours. Strategy 1 provided the largest reduction in sequencing time by sequencing a single sample across two flow cells to reach the needed read requirements. By employing this strategy, a full human genome can be sequenced using 2×100 read lengths at low output mode to >30x coverage within 12 hours (Table 8). When compared to a full length 2×150 high output sequencing run at 31 hours, this is a reduction in run time of 59.7%. Strategy 2 utilized only one flow cell, a single side of the instrument, and made additional adjustments to cycle times to generate data within 24 hours. It utilizes mid output AVITI reagents and generates 500M 2×150 reads in under 24 hours, a reduction of 22.5% compared to a standard 2×150 mid output run. This reduction in sequencing times allows for daily set up and sequencing of samples with minimal reduction in benchmarking performance.

**Table 8:** Rapid WGS run times per sequencing strategy and read length.

| Read Length | Total run time (hr) |  |  |
| --- | --- | --- | --- |
|  | Standard AVITI - 1 FC - CB MO | rWGS Strategy 1 - 2 FC | rWGS Strategy 2 - 1 FC |
| 2x75 | 18.9 | 10.1 | 16.0 |
| 2x100 | 22.9 | 12.5 | 18.6 |
| 2x150 | 31.0 | 17.3 | 24.0 |

While run time was reduced, sequencing quality remained high for both strategies. Strategy 1, in two 12 hour sequencing runs, yielded a mean base quality score >Q40 and >90% Q30 (Figure 9) resulting in marginal increases in error rates during secondary analysis. Using DeepVariant for secondary analysis, total errors increased by 10.08% from 36,207 total benchmarking errors of HG001 in the 2×100 standard Cloudbreak run to 40,267 errors in rWGS-Strategy 1 when analyzed at 30x coverage (Figure 9). rWGS-Strategy 2 yielded similarly high-quality data per single flowcell with a mean base quality score >40 and >92% Q30 quality (Figure 10). Total errors increased marginally, only 3.66% from 32,835 total benchmarking errors of HG002 in the control Cloudbreak FS run to 34,083 total benchmarking errors of HG002 in the rWGS-Strategy 2 sequencing run when down sampled to 35x. Both rWGS workflows balance sequencing speed and throughput by either utilizing both sides of an AVITI instrument concurrently for Strategy 1 (2 flow cells) to generate the requisite read numbers needed to reach 30x coverage of a single human genome (480 M reads @ 2×100 read length) or using a similarly optimized recipe but increasing the total sequencing time to 24 hours on a single side of the instrument yielding 500M, 2×150 read length on a single flow cell.

**Figure 9:**
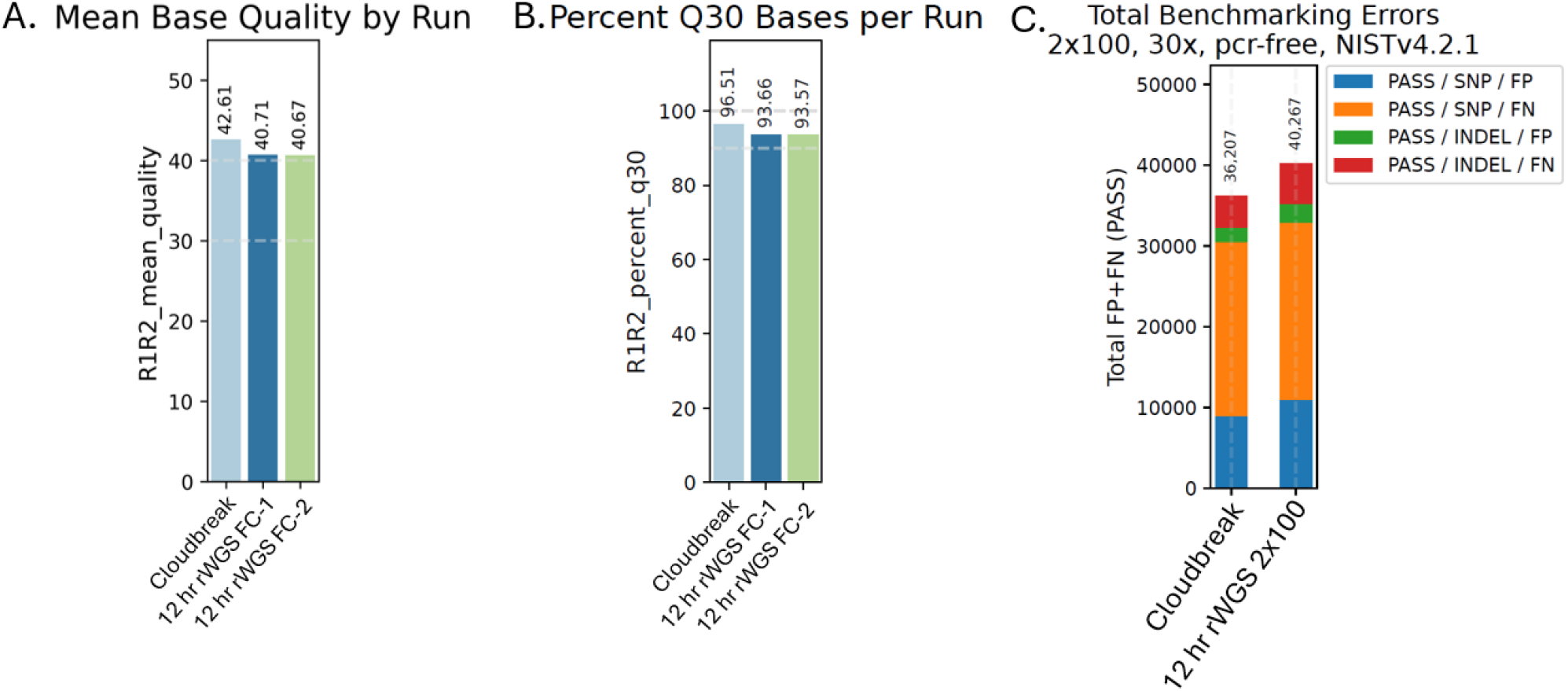
A) Mean base quality by run for each rWGS flow cell sequencing run at 2×100 and standard 2×10 Cloudbreak sequencing run. B) Percent Q30 quality per Run C) Total benchmarking errors for HG001 whe sequenced using either rWGS (40,267) and standard Cloudbreak (36,207) sequencing chemistry. Data was down sampled to 30x using 2×100 read lengths for analysis.

**Figure 10:**
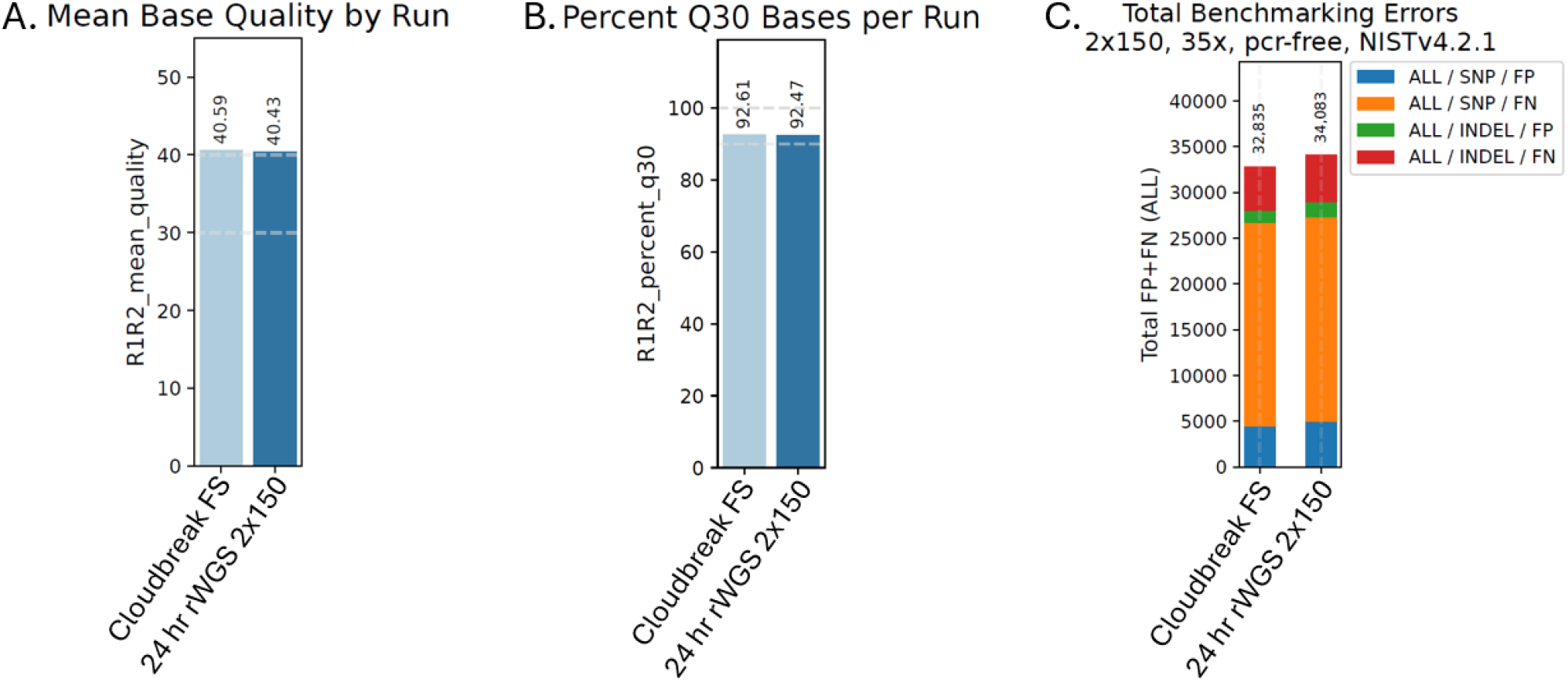
Run metrics for the control Cloudbreak Freestyle (FS) 2×150 and 24hr rWGS 2×150. A) Mean base quality, B) Percent Q30 per run and C) Benchmarking errors of HG002 down sampled to 35x coverage.

## DISCUSSION

In this study, we present approaches that increase analytical precision and accuracy of sequencing data using larger library insert sizes and describe three scenarios of hWGS research projects with various requirements in throughput, ability to map reads through complex regions, and time, all supported by a benchtop sequencing instrument.

### Maximizing Analytical Precision and Accuracy with Larger Library Insert Sizes Under Reduced Coverage Thresholds

WGS and molecular testing is being increasingly used as a front-line screen for patients and is driving discovery of new and unknown variants (Turro et al., 2020). As has been shown through the increase in long-read sequencing, the ability to place reads accurately through repetitive or low-quality contexts increases the information content in your sequencing run, though often coming at an increased cost relative to short-read sequencing. With most existing sequencing platforms on the market having limited capability to accommodate longer insert size (1kb+), library preparation methodologies for short-read sequencers are designed to select libraries to a relatively small size range to maximize sequencing performance. However, rolling circle amplification has fewer biases to smaller fragments (Nallur et al., 2001), and can tolerate these longer sizes. Through the library prep methods introduced in this article in conjunction with sequencing on the AVITI platform the ability to tune and generate libraries with longer median insert sizes and can translate to increased variant calling accuracy (Figure 3).

This effect is seen across all benchmarking regions, including difficult regions (Figure 4A and B), showing the ability of the longer inserts to better anchor reads during alignment and allow for better interrogation of regions of the genome considered “difficult” to analyze. This reduction in SNP FN and improvement in F1 scores have the potential to increase the calling accuracy and precision of rare disease and oncology screening, while keeping the cost and turnaround time benefits of short read sequencing.

### Workflow Optimization Strategies for Effective Sample Handling in Large-Scale Projects

To increase processing throughput and decrease cost per sample, QC assays were developed to miniaturize reagent volume and increase assay plexity. The miniaturized assays utilize 384 well plates and plate readers to multiplex samples across plates and provide confidence in measurements by increasing replicate measures per sample. This miniaturized assay reduces the cost of DNA quantification of 96 samples to $0.36 per sample (n=3 rxn per sample) compared to $0.93 per reaction (n=1 rxn per sample) for the Qubit dsDNA HS assay, resulting in cost reduction per sample by up to 61% and allowing for triplicate reactions to increase accuracy compared to the single-plex standard Qubit dsDNA HS assay.

Efficient and clear tracking of samples is imperative when handling large sample projects and processing samples across multiple sequencing runs over several months. Developing a consistent and clear nomenclature for sample tracking can reduce sample handling errors, provides a link across multiple steps to ensure tracking across workflows, and links final sequencing data to library metadata. Transferring genomic DNA material into Matrix tubes in 96-well plate format allows for individual sample tube barcoding that can be read on a Matrix tube plate reader for quick sample orientation and sample order confirmation which can be critical for sample batching or rearrangement throughout the workflow. As the sample moves through the workflow, new unique IDs are assigned to the sample containing a consistent project suffix, followed by a workflow step classifier then sample number (i.e.: Project Number_Workflow-Sample Number).

### Harnessing Low-Pass ‘Bulk QC’ Sequencing Runs for Improved Utility and Insights

Completing an initial QC sequencing run to measure sample quality and balance is standard practice to reduce the need for re-runs of individual samples. Index CVs within trios vary due to input sample quality, making density targets more difficult to achieve, particularly with the increase in library insert size. Density and loading concentrations tended to fluctuate based on the internal trio index CV with higher intra-trio CV leading to more variation in final density and inaccurate loading concentrations. To address this and limit overall sequencing runs, bulk QC runs were implemented to measure true trio index CV for up to 48 individual samples, as well as provide additional sequencing reads that contribute to the total required reads per sample. Data generated from the same library showed high concordance in Ti:Tv ratios and SNP TP calling between runs, allowing for the combining of data across multiple runs for a single sample with confidence (Table 6, Table 7).

The use of 48-plex (16 trio) Bulk QC sequencing runs improved our intra-trio CV (Table 3). While this does add an additional sequencing run, we found the increase in trio balancing outweighed this extra run. Demonstrating this, we simulated a distribution of 1B sequencing reads spread across 3 samples with a 12.27% unbalanced trio CV vs. 4.41% adjusted trio CV over 1000 replicates using the tidymodels package in R (Kuhn and Wickham, 2020). We find that with the unbalanced trio design, one sample would fail to hit 30x coverage (∼305.5M reads) in 57.4% of the simulations, while the re-balanced trio would fail to hit 30x coverage in only 2.6% of the simulations. This reduces the number of re-queue runs per sample, while also providing initial spot-checks for other sequencing and library metrics.

While many of the samples were high quality gDNA, degradation or contamination from sample collection or extraction caused occasional library prep failures across the batches of library preps. Degraded samples, i.e. gDNA with a low DIN score (<7), yielded smaller linear libraries than samples with larger DIN values (>7). Via sequencing, the gDNA quality variability can be assessed using genomic coverage evenness at 1x from the Bulk QC run data. In this project, we found 42 samples that showed increased genomic variability and non-uniform coverage (CV greater than 0.1 per 100kb coverage window). When investigating the root cause of the increased genomic CV, we found a majority of the libraries with non-uniform coverage were prepared from DNA samples possessing lower gDNA quality, indicating the input gDNA quality is important to achieve even coverage of WGS. This increase in coverage variability can complicate downstream analysis and the ability to confidently call copy number variants (CNVs) above the background coverage level (Trost et al., 2018).

Additionally, through these Bulk QC runs we found that contamination during sample extraction was present in some saliva extracted gDNA, a known issue with this sampling approach (Chrisman et al., 2022; Samson et al., 2020). This contaminated material yields libraries which were indistinguishable from “true” human libraries during initial QC (i.e. normal yields, library sizes), but had low aligned coverage and insert sizes when aligned to human reference. Leveraging the Bulk QC run data prior to full-depth sequencing, we used a metagenomic aligner Kraken 2 (Wood et al., 2019) to screen these low coverage samples, and in a few cases were able to determine a high proportion of reads were from non-human DNA (Figure 11), and the sample would need to be recollected.

**Figure 11:**
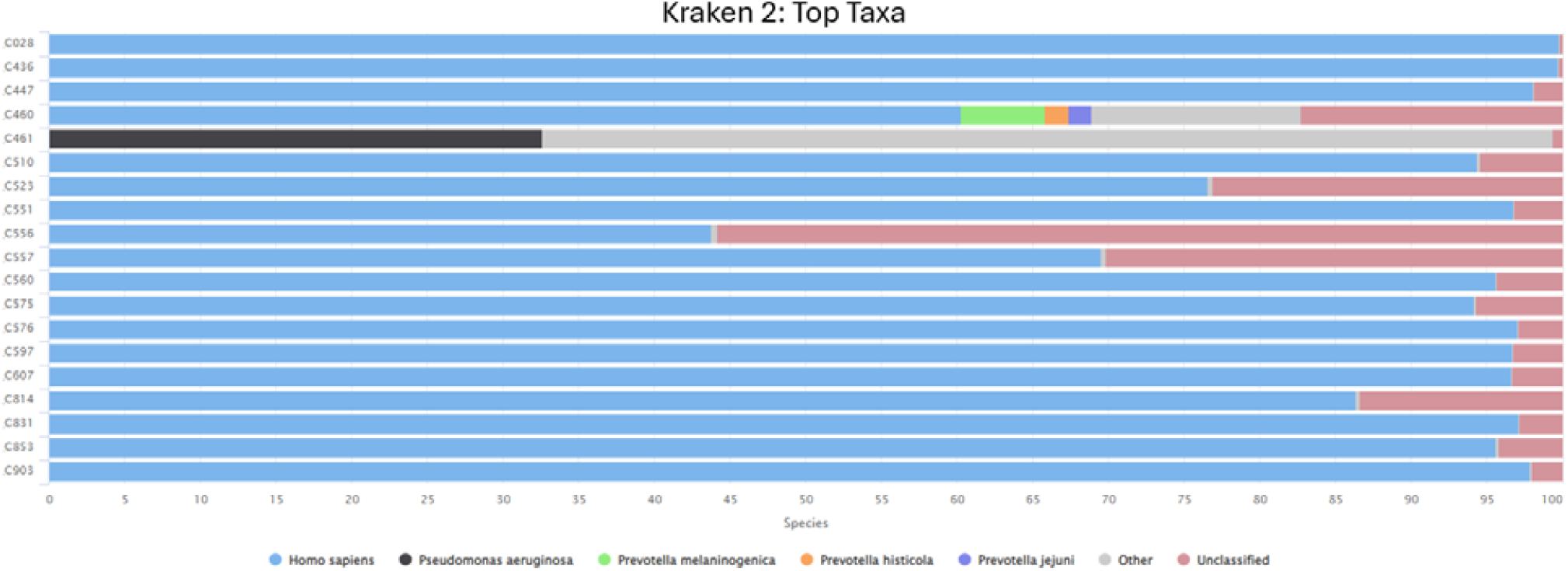
Kraken analysis of human samples potentially contaminated with bacterial species.

Total pass filter polonies per run were tracked over time (Figure 7). After switching library preparation conditions to increase insert size from ∼350 to 600bp, we had a period of lower densities until library loading conditions could be optimized (dashed line 1 in Figure 7a). This relationship of library size and loading concentration is well understood, though should be monitored through a large scale project to maximize throughput. An additional benefit to the bulk QC run methodology is that in addition to the above described library QC data and trio adjustment, you are also able to adjust the overall loading of each trio based on the 48-plex performance due to the consistent performance of the trios between this initial and full density run. Implementation of the Bulk QC runs prior to individual trio sequencing runs improved the output consistency (dashed line 2, Figure 7a), reducing the need for additional top off runs to achieve read depth.

Costs of next generation sequencing are decreasing rapidly, though primarily for the largest scale sequencers. The data and workflows outlined above provide a flexible strategy to scale up sequencing with a modest CapEx budget. With three instruments running concurrently, this strategy would allow for ∼ 2800 human genomes/year with and high flexibility. Not only does this system provide the ability to process thousands of genomes a year at the industry standard coverage, the AVITI system paired with library prep optimization and can generate high quality long insert (>1kb) data capable of elucidating difficult to map regions of the genome and reduce total errors for consistent and accurate variant calling imperative for rare disease and oncology sequencing research applications. Sequencing time can be reduced to ∼12 hours using the rWGS run settings resulting in overall TAT to be drastically shorted where speed is paramount. ABC on AVITI provides a high quality, rapid, flexible, and cost-effective system to reach high throughput sequencing in a small-scale laboratory without the need for sample batching or possess factory scale instrumentation.

